# On a mathematical connection between single-elimination sports tournaments and evolutionary trees

**DOI:** 10.1101/2022.08.09.503313

**Authors:** Matthew C. King, Noah A. Rosenberg

## Abstract

How many ways are there to arrange the sequence of games in a single-elimination sports tournament? We consider the connection between this enumeration problem and the enumeration of “labeled histories,” or sequences of asynchronous branching events, in mathematical phylogenetics. The possibility of playing multiple games simultaneously in different arenas suggests an extension of the enumeration of labeled histories to scenarios in which multiple branching events occur simultaneously. We provide a recursive result enumerating game sequences and labeled histories in which simultaneity is allowed. For a March Madness basketball tournament of 68 labeled teams, the number of possible sequences of games is ~ 1.91 × 10^78^ if arbitrarily many arenas are available, but only ~ 3.60 × 10^68^ if all games must be played sequentially on the same arena.

## 1 A scheduling problem

The National Collegiate Athletic Association men’s and women’s basketball tournaments, colloquially known as “March Madness” after the month during which most of their games take place, are single-elimination sports tournaments with 68 teams from colleges across the United States. Each team is assigned an initial opponent, with subsequent opponents determined by the outcomes of a sequence of specified games. A team that loses a game plays no subsequent games, so that 67 games are played until a single winning team remains.

In typical years, the games are played in multiple distant locations. Games are scheduled in many arenas, often concurrently. The teams in a tournament are divided into four regional groups of approximately equal size (18, 18, 16, and 16 in the 2019 men’s tournament, for example), and they play their games within the regional groups until four teams remain, one from each region. The “Final Four” teams play the last three games in a single arena, revealing the champion of the tournament.

The 2020 tournaments were canceled due to the COVID-19 pandemic. For the 2021 tournaments, with the pandemic continuing, the organizers sought to limit teams’ travel, arranging for all the games to be played in Indiana in the men’s tournament, and San Antonio in the women’s tournament. This circumstance inspires a question:

*Suppose all the games in a single-elimination sports tournament are played sequentially in the same arena. In how many possible sequences can the games be played?*

In an actual March Madness tournament, the sequence of events is divided into “rounds.” First, the “First Four” round is played, reducing the 68 teams to a fully symmetric arrangement of 64 teams. Next, for each *i* from 1 to *k —* 1, with *k* = 6, each remaining team plays its *i*th game before any team plays its (*i* + 1)th game. However, an equally valid sequence would play all the games in one of the four disjoint regional groups of teams before any game is played in the other three groups; such a sequence would identify one member of the “Final Four” before the teams in other regions have played any games at all. Many different sequences of games leading to the “Final Four” can be envisioned.

We will see that the question of enumerating sequences of games reveals connections between a familiar sporting event and problems of evolutionary biology. Although the actual 2021 tournaments used multiple arenas rather than a single one (all in Indiana in the men’s tournament, all in San Antonio in the women’s), the difference from a typical year—with games spread across 14 arenas in 11 states plus the District of Columbia in the 2019 men’s tournament, for example—suffices to inspire a mathematical connection.

## 2 Symmetric single-elimination tournaments

We make the enumeration problem precise. For now, ignore the “First Four” games, and consider a symmetric arrangement of 2^*k*^ distinguishable teams, each of which has the potential to win the tournament by winning k games. Assign each team its sequence of potential opponents. Next, bijectively assign each of the 2^*k*^ – 1 scheduled games a label from {1, 2,..., 2^*k*^ – 1}, subject to a constraint. In particular, a game can only be played after its teams have been determined: the label for a game must exceed the labels for games that determine its teams. We term the tree whose leaves encode the teams and whose interior nodes encode the games a *bracket* (Figure 1).

**Figure 1:**
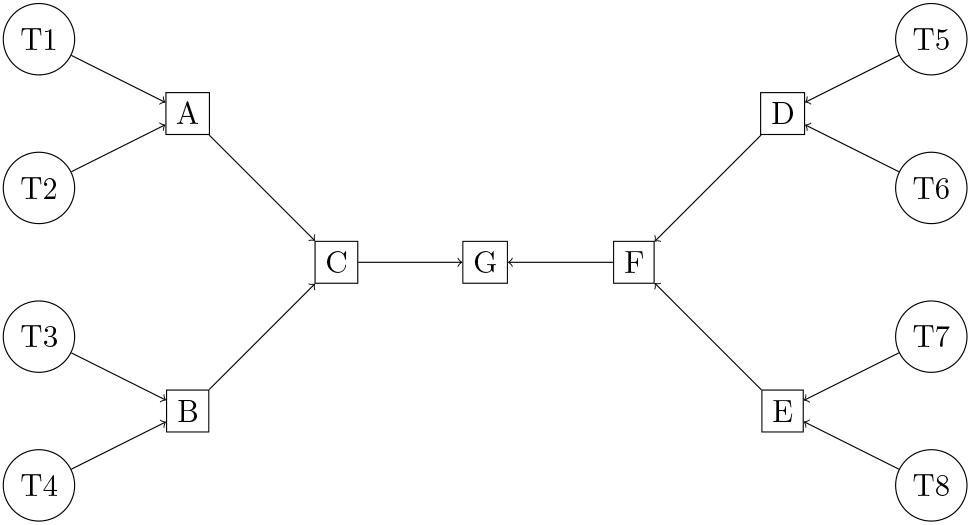
A bracket with 2^3^ = 8 teams. Teams labeled T1 to T8 appear as circles, and games appear as squares. The number of games played is 2^3^ — 1 = 7.

First, we consider a scenario with a single arena. In how many possible sequences can the 2^*k*^ — 1 games of a bracket be played in a single arena? The problem consists in counting permissible bijections between the 2^*k*^ — 1 games and the labels {1,2,..., 2^*k*^ — 1}.

To develop an understanding of the problem, we examine small *k* up to *k* = 6, corresponding to the March Madness men’s tournament from 1985 through 2000 and the women’s tournament from 1994 through 2021; the number of men’s teams was increased from 64 to 65 in 2001 and then 68 in 2011, and the number of women’s teams was increased from 64 to 68 in 2022. For *k* =1, there is a unique ordering of one game. For *k* = 2, either of the two non-championship games can be played first, then the other, and then the championship game—so that there are 2 sequences.

For *k* = 3, consider the bracket diagram in Figure 1. First, note that any number of games among {A, B, C} in the “left” sub-bracket can be played before the first game in the “right” sub-bracket (which is necessarily D or E). Having fixed the schedule of games in each sub-bracket, we can count the number of ways to sequence the games from the two sub-brackets to form a complete sequence of all games; moreover, this number is the same irrespective of the fixed orders of games within sub-brackets. For illustration, suppose the order of games for the left sub-bracket is B, then A, then C. For the right sub-bracket, suppose the order is D, then E, then F.

For fixed sequences of games in the left and right sub-brackets, the number of ways of sequencing the games from the right sub-bracket in relation to the games from the left sub-bracket can be counted by choosing which 3 ranks among {1, 2, 3,4, 5, 6} are assigned to games in the left sub-bracket. For example, suppose the ranks 1, 2,6 are assigned to the left sub-bracket. With the relative game orders B, A, C and D, E, F already chosen, the complete game schedule is B, A, D, E, F, C, followed by G, which must be played last.

Hence, 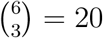 valid game sequences exist for each pair of choices for the order of games in the left and right sub-brackets of 2^2^ = 4 teams. We have already seen, for *k* = 2, that 2 valid sequences of games exist for each of these sub-brackets. Multiplying the number of ways of sequencing the games in the left and right sub-brackets in relation to each other by the numbers of sequences of games in the left and right sub-brackets themselves, the number of possible sequences of games is 20 × 2 × 2 = 80.

This example provides a recurrence. Let *S*(*k*) denote the number of valid game sequences for a tournament of 2^*k*^ teams. We saw *S*(1) = 1, *S*(2) = 2, and *S*(3) = 80 via 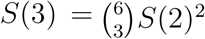. For 2^*k*^ teams, there are *S*(*k* – 1)^2^ pairs of sequences for the two sub-brackets that produce the two teams in the championship game. The binomial coefficient that counts the number of ways that the games of one sub-bracket can be placed in relation to the games of the other sub-bracket is 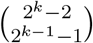. Hence, we have

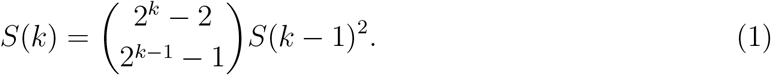

This recurrence, with *S*(1) = 1, has solution

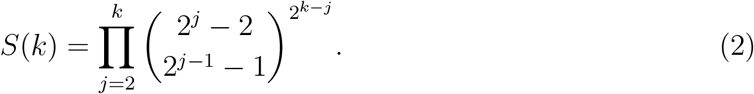

Using eq. 1 or eq. 2, we find

*S*(4) = 21,964,800
*S*(5) = 74,836,825, 861, 835, 980, 800, 000
*S*(6) = 2,606,654, 998,899, 867,556,195, 703, 676, 289, 609, 067, 340, 669, 424, 836, 280, 320, 000, 000, 000.

By a nice coincidence, the quantity *S*(6), counting valid sequences of games in a tournament of 64 teams, has 64 digits.

Note that no particular requirement exists that a bracket has 2^*k*^ games, and the problem of counting possible sequences of games in a bracket can be considered for arbitrary single-elimination tournament structures.

## 3 Labeled histories in evolutionary biology

In fact, the problem of counting sequences of games in a bracket is equivalent to a problem of evolutionary biology, that of enumerating the *labeled histories* that are compatible with a *labeled topology*.

In evolutionary biology, species are related to each other by descent from a common ancestor. The ancestor-descendant relationships can be represented by a tree structure. Consider a rooted binary tree *T* with *y* leaves, bijectively labeled by the elements of a label set containing y elements. In the context of evolution, each label represents the label for a “taxon,” or a distinctive biological group such as a species. The tree describes the descent relationships among the taxa. The rooted labeled binary tree is termed a *labeled topology*. With the precise definition of a bracket above, a bracket is exactly a labeled topology, replacing the leaf labels for biological taxa with the team names in the tournament.

A second structure of interest in evolutionary biology is that of a labeled history. Consider a rooted binary tree *T* as a directed graph with edges that point from the root toward the leaves. A node *w* of *T* is said to be ancestral to a node *v* if *w* lies on the path from the root to *v; v* is said to be descended from *w*. Trivially, a node is both ancestral to and descended from itself. Given a labeled topology for a rooted binary tree *T* with *y* leaves, a *labeled history* for *T* is a bijection *σ* from {1, 2,..., *y* – 1} to the internal (non-leaf) nodes of *T*, satisfying the constraint that if node *v* is descended from node *w* in *T*, then *σ*^-1^(*v*) ≤ *σ*^-1^ (*w*).

Observe that this constraint on labeled histories is precisely the constraint that makes a sequence of games valid for a tournament: valid game sequences compatible with a tournament correspond to labeled histories compatible with a labeled topology. Hence, we can use ideas from the mathematical study of evolutionary trees, or *mathematical phylogenetics*, for sports tournaments, and vice versa.

Note that in an evolutionary tree, the branching of one lineage into two is treated as taking place instantaneously. In the analogy with sports tournaments, it is convenient to also assume that each game is played instantaneously. Note also that for alignment with the sports analogy, in our definition of labeled histories here, the numbering of internal nodes increases from leaves toward the root, but in mathematical-phylogenetic studies, the opposite convention is often adopted.

### 3.1 Review of mathematical phylogenetics results

The problem of enumerating labeled histories compatible with a labeled topology can be solved recursively [15]. Consider a labeled topology *T* with subtrees *L* and *R* immediately descended from the root, where one subtree (*L*) is arbitrarily assigned to be the “left” subtree and the other (*R*) is arbitrarily assigned to be the “right” subtree. Let |*T*| be the number of leaves of *T*, so that *T* has |*T*| – 1 internal nodes. For each pair consisting of a labeled history for *L* and a labeled history for *R*, we must count the number of ways of placing into a sequence the |*L*| – 1 internal nodes of *L* and the |*R*| – 1 internal nodes of *R*. Reserving the label |*T*| – 1 for the root node, |*L*| — 1 of the numbers {1, 2,..., |*T*| – 2} must be assigned to the internal nodes of *L* and the rest to the internal nodes of *R*. This assignment can be made in 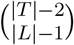 ways. Hence, multiplying by the numbers of labeled histories for *L* and *R*, and noting that the number of labeled histories is *N*(*T*) = 1 for labeled topologies *T* with 1 leaf (and for those with 2 and 3 leaves), we have a recurrence.

#### Theorem 1.

*For a labeled topology T, the number of labeled histories N*(*T*) *is for*

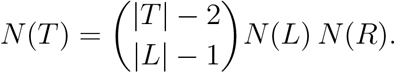

for |*T*| ≥ 2, *with N*(*T*) = 1 *for* |*T*| = 1.

This recurrence can produce a non-recursive formula [5]. For subtrees *L* and *R*, let *their* subtrees be *L_ℓ_, L_r_* and *R_ℓ_*, *R_r_*, respectively. Applying the recurrence,

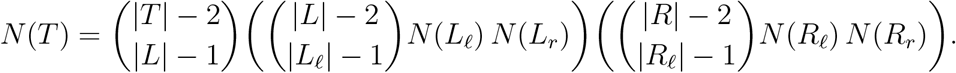

Iterating until each subtree in the expression has 1, 2, or 3 leaves, we have a product of binomial coefficients, one for each internal node of *T*. To state the result precisely, denote by *V*^0^(*T*) the set of internal nodes of *T*, including the root. For each *v* ∈ *V*^0^(*T*), denote by *m*(*v*) the number of leaves in the subtree rooted at *v*, and denote by *ℓ*(*v*) the number of leaves in the left subtree of the subtree rooted at *v*. Then

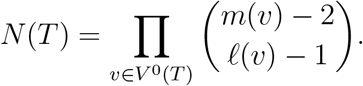

In this product, each internal node other than the root appears in the numerator of one binomial coefficient and the denominator of another; the root appears only in a numerator. Multiplying by (|*T*| – 1)/(|*T*| – 1), the expression can be simplified.

#### Theorem 2.

*For a labeled topology T, the number of labeled histories N*(*T*) *is*

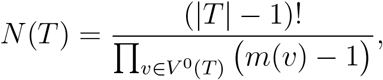

*for* |*T*| ≥ 2, *with N*(*T*) = 1 *for* |*T*| = 1.

As an example of the theorem, consider the case in which *T* is the labeled topology that corresponds to the symmetric bracket for 64 teams (Figure 2). Considering all internal nodes *v* in T, the tree contains one node with 64 descendant leaves, two nodes with 32 descendant leaves, four nodes with 16 descendant leaves, eight nodes with 8 descendant leaves, 16 nodes with 4 descendant leaves, and 32 nodes with 2 descendant leaves. Hence, the product over internal nodes in Theorem 2 is

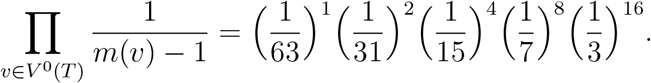

**Figure 2:**
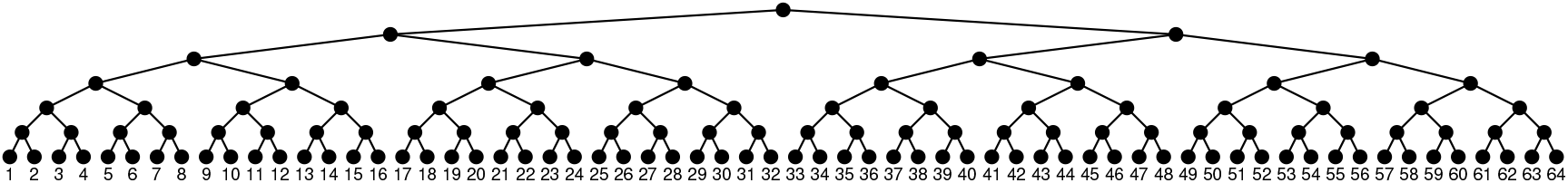
A labeled topology with 64 labeled leaves. This labeled topology corresponds to a 64-team bracket, with teams numbered 1 to 64; the root node is the championship game.

Multiplying by 63! recovers the product from eq. 2, *N*(*T*) = *S*(6) ≈ 2.61 × 10^63^.

The full March Madness men’s tournament bracket from 2021, with the “First Four” games included, adds two games each to two of the four sub-brackets that produce the “Final Four” teams, so that two sub-brackets contain 18 teams and the other two contain 16 teams.

This modification of the symmetric bracket for 64 teams produces 4 more internal nodes (Figure 3). It also changes the numbers of descendants for many of the nodes in the resulting tree *T*. Considering all internal nodes *v, m*(*v*) now takes values 68, 36, 32, 18, 16, 9, 8, 5, 4, 3, and 2. The numbers of nodes with these numbers of descendants are 1, 1, 1, 2, 2, 4, 4, 4, 12, 4, and 34, respectively. The product in Theorem 2 becomes

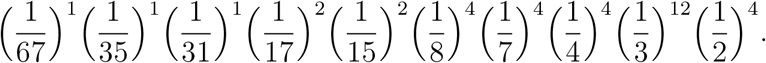

**Figure 3:**
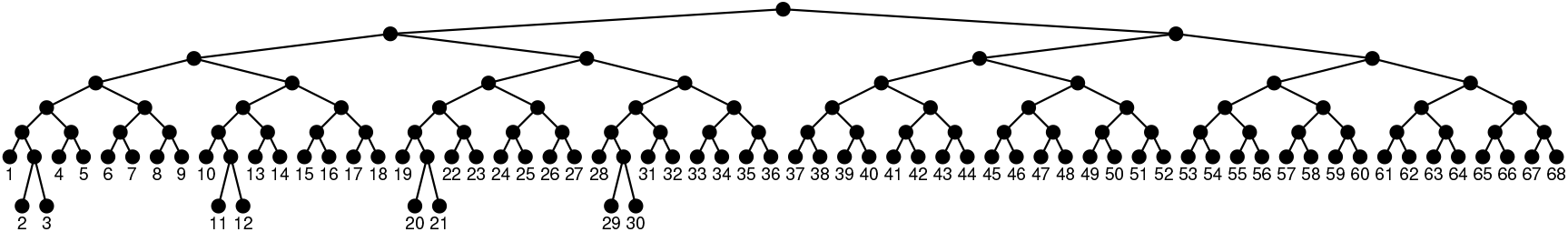
A labeled topology with 68 labeled leaves. This labeled topology corresponds to the 68-team bracket that was used in the March Madness men’s tournament in 2021.

Multiplying by 67! gives *N*(*T*) ≈ 3.60 × 10^68^ possible sequences of games.

### 3.2 Uses of labeled histories

Labeled histories, sometimes termed *coalescence sequences* or *ranked labeled trees*, appear frequently in mathematical phylogenetics. They are among the main classes of tree structures used in assessing probabilistic outcomes of assumptions about evolution [28, p. 47]. It is often convenient for evolutionary models to assume that each sequence of branching events that could produce a rooted binary tree for a set of labeled species is equally likely; this assumption, that of the *Yule* or *Yule–Harding* model in phylogenetics [1, 13, 15, 20, 27, 28, 30, 33, 35], produces a uniform distribution on labeled histories.

Computations concerning features of tree shape for evolutionary trees often evaluate the probability that such features are produced under the Yule–Harding model, so that they directly or indirectly examine the fraction of labeled histories on y leaves that possess a given feature, or the probability distribution of a quantity across labeled histories [3, 5, 6, 7, 8, 10, 12, 16, 18, 22, 24, 25, 29, 31, 34, 36, 37]. Mathematical phylogenetics computations have used combinatorial results on the set of labeled histories for y species, for example employing a space of labeled histories with a notion of distance between them [26] and a characterization of the labeled topologies that possess the largest number of labeled histories [9, 11, 14]. In some situations, phylogenetic research reports results on labeled histories that have been obtained in equivalent scenarios in computer science, involving concepts such as binary search trees [4, 17, 21], heaps [32, p. 1319], and tableaux [19, p. 67, problem 20].

## 4 Simultaneous games, simultaneous binary mergers

Not only does the biological setting introduce a result for the sports tournament sequence enumeration problem, the sports context introduces a new idea that has not often been considered in the biological setting of evolutionary trees: simultaneity. If the games of a tournament are played on multiple arenas, as is true of March Madness in typical years, then games can be played simultaneously.

Suppose *ℓ* arenas are available. A sequence of games is now permitted to possess “ties,” where a tie represents games played on different arenas at the same time. How many *tie-permitting* sequences are possible for a tournament with bracket *T* if *ℓ* arenas are available?

We call this quantity *N^*ℓ*^* (*T*); *N*^1^(*T*) gives the case with one arena, denoted by *N*(*T*) in Theorems 1 and 2. We enumerate tie-permitting sequences in the “infinite-arenas” setting, obtaining *N*^∞^(*T*). The number of available arenas is not actually infinite, but simply satisfies 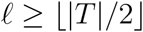, where |*T*| is the number of teams in *T*, as no more than 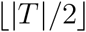 games can ever be played simultaneously in bracket *T*.

For convenience, we refer to a collection of simultaneous games as an “event.” A feature of the infinite-arenas context is that events induced by two game sequences, one for the left sub-bracket of a node and the other for the right sub-bracket, can be combined into a joint event when forming the complete game sequence, without occupying more arenas than are available. By contrast, if, for example, *ℓ* =2, then a 2-game event from the left sub-bracket cannot be simultaneous with an event from the right sub-bracket, as this joint event would occupy at least 3 arenas.

To state a succinct recurrence for *N*^∞^(*T*), we let *E*^∞^(*T, n*) be the number of tie-permitting sequences on the bracket *T* which consist of exactly *n* events, so that the sequence occupies exactly n distinct points in time. Note that *E^∞^*(*T*, |*T*| – 1) is equal to *N^X^*(*T*), or equivalently, the quantity denoted by *N*(*T*) in Theorems 1 and 2. Let *δ*(*T*) denote the depth of *T*, the maximum length of a path from the root of *T* to one of its leaves. We have

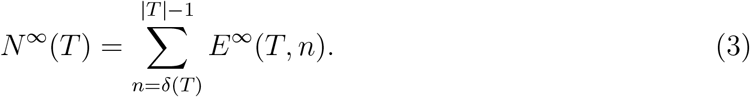

We give the following recurrence for *E*^∞^(*T, n*), noting that for a trivial bracket *T* consisting of a single team, *E*^∞^(*T*, 0) = 1 and *E*^∞^(*T, n*) = 0 for *n* ≠ 0.

### Theorem 3.

*Let T be a bracket with left sub-bracket L and right sub-bracket R.*

i. *If* (|*L*|, |*R*|) = (1,1), *then E*^∞^(*T*, 1) = 1 *and E*^∞^(*T, n*) = 0 *for n* =1.
ii. *If at least one of* |*L*|, |*R*| *exceeds 1, then*

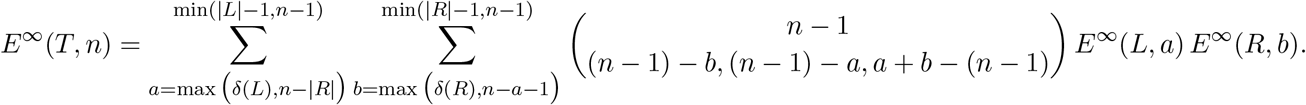

*Proof.* For (i), if a bracket *T* has only one team in the left sub-bracket and one team in the right sub-bracket, then a single game is played (*n* = 1), and trivially, only one sequence exists for this game. Hence, *E*^∞^(*T*, 1) = 1 and *E*^∞^(*T, n*) = 0 for *n* ≠ 1.

For the recursive case (ii), we count sequences that merge a sequence of games from the left sub-bracket *L* and a sequence of games from the right sub-bracket *R*. The last event in any such sequence is a single game, corresponding to the root node of *T*.

In a sequence of *n* events, the number *a* of events in *L* can range from *δ*(*L*) to |*L*| – 1, and the number *b* of events in *R* can range from *δ*(*R*) to |*R*| – 1. We impose the additional conditions that *a, b* ≤ *n* – 1 to ensure that at most *n* – 1 events are used for the bracket *T* and *a* + *b* ≥ *n* – 1 to ensure that *at least n* – 1 events are used.

The number of sequences for the left sub-bracket containing *a* events is *E*^∞^(*L, a*), and the number for the right sub-bracket containing b events is *E*^∞^ (*R,b*). It remains to prove that the trinomial coefficient correctly counts the number of possible ways to form a sequence of *n* – 1 events for the bracket *T* (excluding the root) by combining the a events of the left sub-bracket and the *b* events of the right sub-bracket in an order-preserving, tie-permitting sequence. Each of the *n* – 1 events must be formed in one of three ways: an event among the *a* events of the left sub-bracket occurs and is not simultaneous with an event among the *b* events of the right sub-bracket, an event among the *b* events of the right sub-bracket occurs and is not simultaneous with an event among the *a* events of the left sub-bracket, or an event among the *a* events of the left sub-bracket is simultaneous with an event among the *b* events of the right sub-bracket. The numbers of events in these three disjoint categories are (*n* – 1) – *b*, (*n* – 1) ‒ *a*, and *a* + *b* — (*n* – 1), respectively.

In the biological context, the setting of Theorem 3 corresponds to labeled histories with ties, in which—looking backward in time—multiple pairs of lineages can coalesce simultaneously. How many tie-permitting labeled histories are possible for a labeled topology if arbitrarily many pairwise coalescences can occur simultaneously? In mathematical evolutionary biology, models that permit simultaneous pairwise coalescences, or *simultaneous binary mergers*, are sometimes studied [2, 23]. Such models relax the assumption of the Yule-Harding model for tree shape that coalescences must be asynchronous. They are useful when considering genealogies of genetic lineages sampled in small populations in discrete time; in such settings, it is not improbable that multiple pairs of lineages will coalesce in the same discrete time step. The problem of counting sequences of games for single-elimination tournaments when multiple arenas are available is the problem of counting tie-permitting labeled histories when arbitrarily many simultaneous binary mergers are permissible. Hence, Theorem 3, allowing simultaneous binary mergers in the enumeration of labeled histories, generalizes the recursion of Theorem 2 that counts labeled histories in the standard setting when ties are not permitted.

For tournaments with 2^*k*^ teams, we can compare the number of sequences of games in the arbitrary-arenas case in Theorem 3 to the single-arena case in eq. 2 (Table 1). In the case of 2^2^ = 4 teams, the arbitrary-arenas case permits one additional sequence that cannot occur with only a single arena: a sequence in which two games are played simultaneously in two arenas. For 2^3^ = 8 teams, 365 sequences are possible when arbitrarily many arenas are available, compared to the 80 possible with only one arena.

**Table 1:**
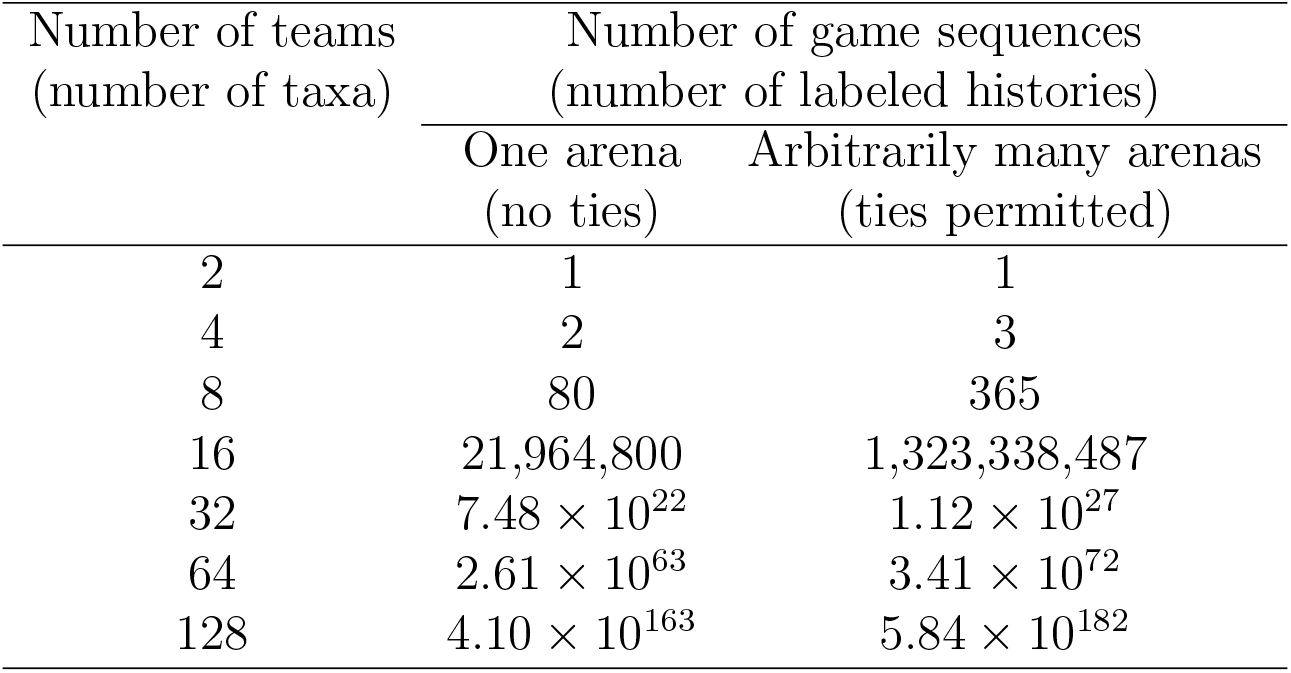
The number of sequences of games for symmetric brackets of 2^*k*^ teams, *k* = 1, 2, 3, 4, 5, 6, 7, with one arena (Theorem 2) or arbitrarily many arenas (Theorem 3) — or, the number of labeled histories for symmetric labeled topologies of 2^*k*^ taxa, either disallowing or allowing ties in coalescence times. For *k* = 5 to *k* = 7, the values are approximate. For the 68-team bracket in Figure 3, the number of game sequences is ~ 3.60 × 10^68^ in one arena and ~ 1.91 × 10^78^ for arbitrarily many arenas.

| Number of teams<br>(number of taxa) | Number of game sequences<br>(number of labeled histories) |  |
| --- | --- | --- |
|  | One arena<br>(no ties) | Arbitrarily many arenas<br>(ties permitted) |
| 2 | 1 | 1 |
| 4 | 2 | 3 |
| 8 | 80 | 365 |
| 16 | 21,964,800 | 1,323,338,487 |
| 32 | $7.48 \times 10^{22}$ | $1.12 \times 10^{27}$ |
| 64 | $2.61 \times 10^{63}$ | $3.41 \times 10^{72}$ |
| 128 | $4.10 \times 10^{163}$ | $5.84 \times 10^{182}$ |

The number of sequences in the arbitrary-arenas case grows rapidly in relation to the number in the single-arena case. For 2^4^ = 16 teams in a symmetric bracket, as used for the single-elimination round of World Cup soccer, the number of sequences is 1, 323, 338, 487 for the arbitrary-arenas case compared to 21, 964, 800 sequences for the single-arena case. For 2^7^ = 128 teams in a symmetric bracket, as used for Grand Slam tennis tournaments, the number is approximately 5.84 × 10^182^ for the arbitrary-arenas case compared to 4.10 × 10^163^ for the single-arena case. For March Madness, the 68-team design in Figure 3 gives 1.91 × 10^78^ sequences compared to 3.60 × 10^68^.

Tables 2 and 3 show the numbers of game sequences for all tournament designs with up to 8 teams — the numbers of tie-permitting labeled histories for all labeled topologies with at most 8 taxa. In these tables, for a tree *T*, the number of sequences for the single-arena case can be seen in the column for *n* = |*T*| – 1 events, as each of the |*T*| – 1 games represents a distinct event.

**Table 2:** The number of sequences of games for brackets with at most 7 teams. For convenience, each bracket *T* is depicted as unlabeled, so that the leaf labeling is omitted. The entries represent the terms *E*^∞^(*T, n*) in Theorem 3, with sum *N*^∞^(*T*) (eq. 3).

| Number of teams<br>(number of taxa) | Bracket<br>(labeled<br>(topology)) | Number of game sequences with $n$ events<br>(number of labeled histories with $n$ events) | | | | | | |
| --- | --- | --- | --- | --- | --- | --- | --- | --- |
| | | $n = 1$ | $n = 2$ | $n = 3$ | $n = 4$ | $n = 5$ | $n = 6$ | Total |
| 2 |  | 1 |  |  |  |  |  | 1 |
| 3 |  |  | 1 |  |  |  |  | 1 |
| 4 |  |  |  | 1 |  |  |  | 1 |
| 4 |  |  | 1 | 2 |  |  |  | 3 |
| 5 |  |  |  |  | 1 |  |  | 1 |
| 5 |  |  |  | 1 | 2 |  |  | 3 |
| 5 |  |  |  | 2 | 3 |  |  | 5 |
| 6 |  |  |  |  |  | 1 |  | 1 |
| 6 |  |  |  |  | 1 | 2 |  | 3 |
| 6 |  |  |  |  | 2 | 3 |  | 5 |
| 6 |  |  |  |  | 3 | 4 |  | 7 |
| 6 |  |  |  | 2 | 9 | 8 |  | 19 |
| 6 |  |  |  | 1 | 6 | 6 |  | 13 |
| 7 |  |  |  |  |  |  | 1 | 1 |
| 7 |  |  |  |  |  | 1 | 2 | 3 |
| 7 |  |  |  |  |  | 2 | 3 | 5 |
| 7 |  |  |  |  |  | 3 | 4 | 7 |
| 7 |  |  |  |  | 2 | 9 | 8 | 19 |
| 7 |  |  |  |  | 1 | 6 | 6 | 13 |
| 7 |  |  |  |  |  | 4 | 5 | 9 |
| 7 |  |  |  |  | 3 | 12 | 10 | 25 |
| 7 |  |  |  |  | 6 | 20 | 15 | 41 |
| 7 |  |  |  |  | 3 | 12 | 10 | 25 |
| 7 |  |  |  | 1 | 12 | 30 | 20 | 63 |

**Table 3:**
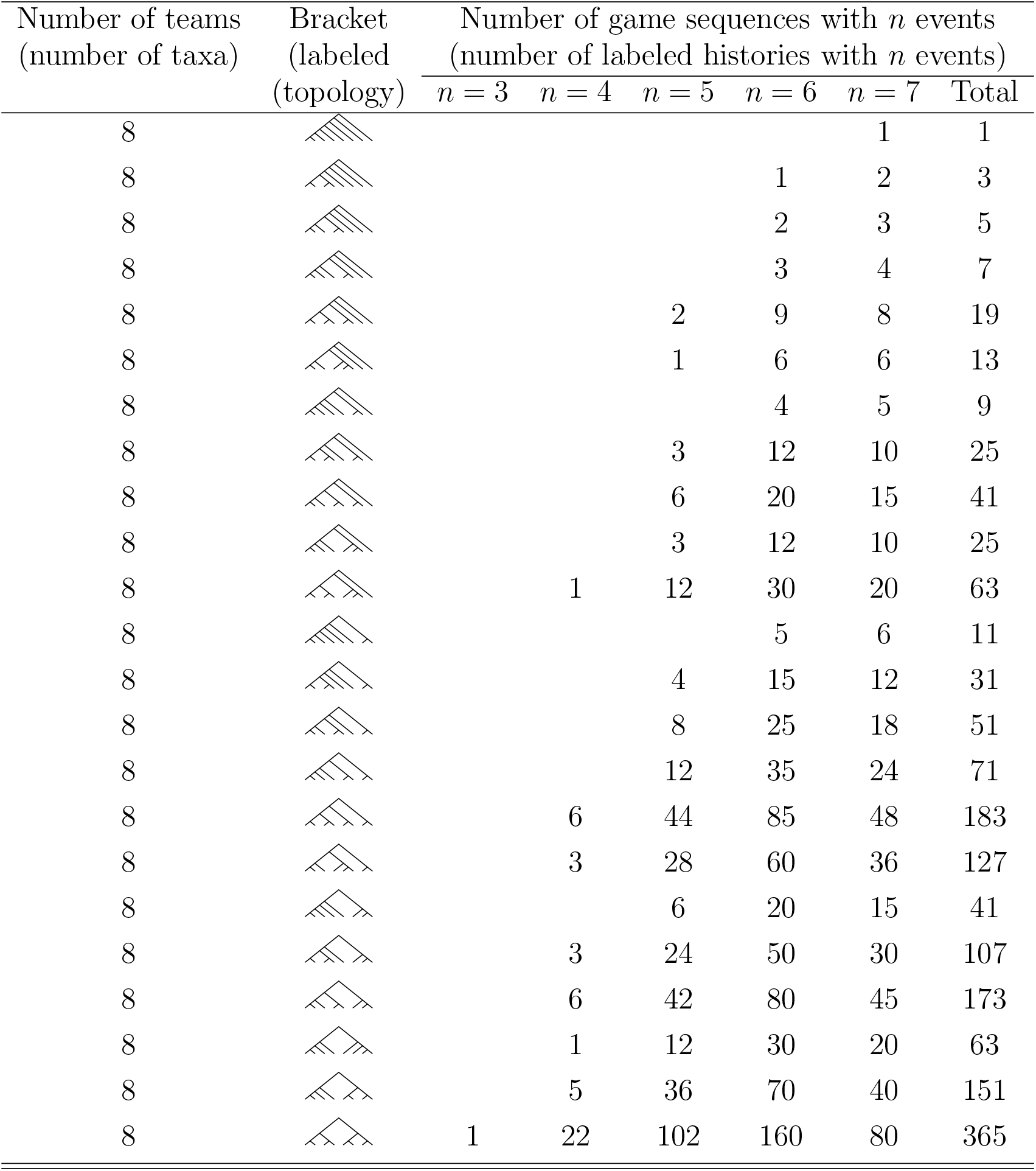
The number of sequences of games for brackets with 8 teams. For convenience, each bracket *T* is depicted as unlabeled, so that the leaf labeling is omitted. The entries represent the terms *E*^∞^(*T,n*) in Theorem 3, with sum *N*^∞^(*T*) (eq. 3).

In Tables 2 and 3, for fixed trees, we can compare the numbers of sequences with different numbers of events. The number of sequences tends to increase as the number of events increases from its minimum, *δ*(*T*), reaching a peak before decreasing as the number of events reaches its maximum, |*T*| – 1. For example, for the fully symmetric bracket with 8 teams, the number of sequences increases from 1 with 3 events to 22 with 4 events, 102 with 5 events, and 160 with 6 events, before declining to 80 with 7 events. The minimal number of events, *δ*(*T*), introduces a constraint that requires many specific games to be played simultaneously; somewhat larger values for the number of events are less constrained, allowing larger numbers of game sequences. It will be of interest to more formally explore this pattern of change with *n* for fixed *T*.

For a specified number of teams, considering different trees, the number of tie-permitting sequences tends to increase with an increasing amount of “balance” in the tree structure. Thus, for example, with 8 teams, the “caterpillar” bracket in the first row of Table 3 possesses only one sequence, whereas the fully symmetric bracket in the last row possesses the largest number, 365. This observation, that the number of tie-permitting sequences increases with tree balance, considered informally, follows a pattern seen in the single-arena case [14]. It suggests the possibility of searching for results concerning the tree shapes that produce the largest number of labeled histories when ties are permitted.

## 5 Conclusion

We have illuminated a connection between labeled histories in phylogenetics and sequences in which the games of a single-elimination tournament can be played on a single arena. By also obtaining a recursion that counts the number of sequences of games for single-elimination tournaments when arbitrarily many arenas are available, we have identified the equivalent problem of counting tie-permitting labeled histories when simultaneous coalescence events are permitted, and we have provided a recursive solution.

On April 4, 2021, Stanford defeated Arizona 54-53 at the Alamodome in San Antonio in the last coalescence of the 2021 March Madness women’s tournament. The following day, Baylor defeated Gonzaga 86-70 at Lucas Oil Stadium in Indianapolis in the corresponding last coalescence for the men’s tournament. Interestingly, the asynchronicity of these championship games, enabling audiences to watch both of them, suggests a more general question of counting game sequences that either asynchronously interleave the games of multiple tournaments or that permit synchronous games across tournaments: the question of counting labeled histories, without and with simultaneous binary mergers, for forests of labeled topologies.

An amusing problem concerning a resourceful attempt to conduct sporting events under the constraints induced by the COVID-19 pandemic has revealed new results for combinatorial structures in evolutionary biology. And if the schedulers of March Madness ever find themselves in circumstances that demand that all the games in a 68-team tournament (as in Figure 3) must be played in one arena, they can rest assured that although the number of available game sequences for a specified bracket is greatly reduced from the 1,905,458,855,466,636,787,971,925,146,177,334,793,473,753,765,414,856,950,607,419,556,152,726,849,614,067 that are permissible in the case that arbitrarily many arenas are available, the single-arena scenario still leaves them with 360,410,120,625,822,474,490,741,822,944,015,962,624,736,196,480,481,624,064,000,000,000,000 possibilities.

## Acknowledgments

We acknowledge NSF grant BCS-2116322 for support.

## References

[1] Aldous, D. J. (2001) Stochastic models and descriptive statistics for phylogenetic trees, from Yule to today. Stat. Sci. 16: 23–34. doi.org/10.1214/ss/998929474

[2] Bhaskar, A., Clark, A. G., Song. Y. S. (2014). Distortion of genealogical properties when the sample is large. Proc. Natl. Acad. Sci. USA, 111: 2385–2390. doi.org/10.1073/pnas.1322709111

[3] Blum, M. G. B., François, O. (2005). On statistical tests of phylogenetic tree imbalance: the Sackin and other indices revisited. Math. Biosci. 195: 141–153. doi.org/10.1016/j.mbs.2005.03.003

[4] Blum, M. G. B., Françcois, O., Janson, S. (2006). The mean, variance and limiting distribution of two statistics sensitive to phylogenetic tree balance. Adv. Appl. Prob. 16: 2195–2214. doi.org/10.1214/105051606000000547

[5] Brown, J. K. M. (1994). Probabilities of evolutionary trees. Syst. Biol. 43: 78–91, 1994. doi.org/10.1093/sysbio/43.1.78

[6] Chang, H., Fuchs, M. (2010). Limit theorems for patterns in phylogenetic trees. J. Math. Biol. 60: 481–512. doi.org/10.1007/s00285-009-0275-6

[7] Choi, K. P., Thompson, A., Wu, T. (2020). On cherry and pitchfork distributions of random rooted and unrooted phylogenetic trees. Theor. Pop. Biol. 132: 92–104. doi.org/10.1016/j.tpb.2020.02.001

[8] Choi, K. P., Kaur, G., Wu., T. (2021). On asymptotic joint distributions of cherries and pitchforks for random phylogenetic trees. J. Math. Biol., 83: 40. doi.org/10.1007/s00285-021-01667-2

[9] Degnan, J. H., Rosenberg, N. A. (2006). Discordance of species trees with their most likely gene trees. PLoS Genet. 2: 762–768. doi.org/10.1371/journal.pgen.0020068

[10] Disanto, F., Wiehe, T. (2013). Exact enumeration of cherries and pitchforks in ranked trees under the coalescent model. Math. Biosci. 242: 195–200. doi.org/10.1016/j.mbs.2013.01.010

[11] Disanto, F., Rosenberg, N. A. (2017). Enumeration of ancestral configurations for matching gene trees and species trees. J. Comput. Biol. 24: 831–850. doi.org/10.1089/cmb.2016.0159

[12] Disanto, F., Fuchs, M., Pangingbatan, A. R., Rosenberg, N. A. (2022). The distributions under two species-tree models of the number of root ancestral configurations for matching gene trees and species trees. Ann. Appl. Prob. 32: 10.1214/22-AAP1791.

[13] Edwards, A. W. F. (1970). Estimation of the branch points of a branching diffusion process. J. Roy. Statist. Soc. Ser. B 32: 155–174. doi.org/10.1111/j.2517-6161.1970.tb00828.x

[14] Hammersley, J. M., Grimmett, G. R. (1974). Maximal solutions of the generalized subadditive inequality. In: Harding, E. F., Kendall, D. G., eds. Stochastic Geometry. London: Wiley, pp. 270–285.

[15] Harding, E. F. (1971). The probabilities of rooted tree-shapes generated by random bifurcation. Adv. Appl. Prob. 3: 44–77. doi.org/10.2307/1426329

[16] Heard, S. B. (1992). Patterns in tree balance among cladistic, phenetic, and randomly generated phylogenetic trees. Evolution 46: 1818–1826. doi.org/10.1111/j.1558-5646.1992.tb01171.x

[17] Holmgren, C., Janson, S. (2015). Limit laws for functions of fringe trees for binary search trees and random recursive trees. Electron. J. Probab. 20: 1–51. doi.org/10.1214/EJP.v20-3627

[18] Kirkpatrick, M., Slatkin, M. (1993). Searching for evolutionary patterns in the shape of a phylogenetic tree. Evolution 47: 1171–1181. doi.org/10.1111/j.1558-5646.1993.tb02144.x

[19] Knuth, D. E. (1998). The Art of Computer Programming Volume 3, 2nd ed. Reading, MA: Addison-Wesley.

[20] Lambert, A., Stadler, T. (2013). Birth-death models and coalescent point processes: The shape and probability of reconstructed phylogenies. Theor. Pop. Biol. 90: 113–128. doi.org/10.1016/j.tpb.2013.10.002

[21] Mahmoud, H. M. (1992). Evolution of Random Search Trees. New York: Wiley.

[22] McKenzie, A., Steel, M. (2000). Distributions of cherries for two models of trees. Math. Biosci. 164: 81–92. doi.org/10.1016/s0025-5564(99)00060-7

[23] Melfi, A., Viswanath, D. (2018). Single and simultaneous binary mergers in Wright–Fisher genealogies. Theor. Pop. Biol. 121: 60–71. doi.org/10.1016/j.tpb.2018.04.001

[24] Rosenberg, N. A. (2003). The shapes of neutral gene genealogies in two species: probabilities of monophyly, paraphyly, and polyphyly in a coalescent model. Evolution 57: 1465–1477. doi.org/10.1111/j.0014-3820.2003.tb00355.x

[25] Rosenberg, N. A. (2006). The mean and variance of the numbers of r-pronged nodes and r-caterpillars in Yule-generated genealogical trees. Ann. Combinator. 10: 129–146. doi.org/10.1007/s00026-006-0278-6

[26] Song, Y. S. (2006). Properties of subtree-prune-and-regraft operations on totally-ordered phylogenetic trees. Ann. Combinator. 10: 147–163. doi.org/10.1007/s00026-006-0279-5

[27] Steel, M. (2014). Tracing evolutionary links between species. Amer. Math. Monthly 121: 771–792. doi.org/10.4169/amer.math.monthly.121.09.771

[28] Steel, M. (2016). Phylogeny: Discrete and Random Processes in Evolution. Philadelphia: Society for Industrial and Applied Mathematics.

[29] Steel, M., McKenzie, A. (2001). Properties of phylogenetic trees generated by Yule-type speciation models. Math. Biosci. 170: 91–112. doi.org/10.1016/s0025-5564(00)00061-4

[30] Steel, M., McKenzie, A. (2002). The ‘shape’ of phylogenies under simple random speciation models. In: Lässig, M., Valleriani, A., eds. Biological Evolution and Statistical Physics. Berlin: Springer, pp. 162–180.

[31] Than, C. V., Rosenberg, N. A. (2014). Mean deep coalescence cost under exchangeable probability distributions. Discr. Appl. Math. 174: 11–26. doi.org/10.1016/j.dam.2014.02.010

[32] Weisstein, E. (2002). CRC Concise Encyclopedia of Mathematics, 2nd ed. Boca Ration: CRC Press.

[33] Wiehe, T. (2021). Counting, grafting and evolving binary trees. In: Baake, E., Wakolbinger, A., eds. Probabilistic Structures in Evolution. Zurich: EMS Publishing House, pp. 427–450.

[34] Wu, T., Choi, K. P. (2016). On joint subtree distributions under two evolutionary models. Theor. Pop. Biol. 108: 13–23. doi.org/10.1016/j.tpb.2015.11.004

[35] Yule, G. U. (1925). A mathematical theory of evolution based on the conclusions of Dr. J. C. Willis, F. R. S. Phil. Trans. R. Soc. Lond. B 213: 21–87. doi.org/10.1098/rstb.1925.0002

[36] Zhu, S., Degnan, J. H., Steel, M. (2011). Clades, clans, and reciprocal monophyly under neutral evolutionary models. Theor. Pop. Biol. 79: 220–227. doi.org/10.1016/j.tpb.2011.03.002

[37] Zhu, S., Than, C., Wu, T. (2015). Clades and clans: a comparison study of two evolutionary models. J. Math. Biol. 71: 99–124. doi.org/10.1007/s00285-014-0817-4

